# Streamline Density Normalization: A Robust Approach to Mitigate Bundle Variability in Multi-Site Diffusion MRI

**DOI:** 10.1101/2025.08.18.670965

**Authors:** Yixue Feng, Yuhan Shuai, Julio E. Villalón-Reina, Bramsh Q. Chandio, Sophia I. Thomopoulos, Talia M. Nir, Neda Jahanshad, Paul M. Thompson

## Abstract

Tractometry enables quantitative analysis of tissue microstructure is sensitive to variability introduced during tractography and bundle segmentation. Differences in processing parameters and bundle geometry can lead to inconsistent streamline reconstructions and sampling, ultimately affecting the reproducibility of tractometry analysis. In this study, we introduce Streamline Density Normalization (SDNorm), a supervised two-step method designed to reduce variability in bundle reconstructions. SDNorm first computes streamline weights using linear regression to match a subject’s bundle to a template streamline density map, then iteratively prunes streamlines to achieve a target density using a novel metric called *effective Streamline Point Density* (eSPD). We evaluate SDNorm across multiple bundles and acquisition protocols in dMRI data from a subset of subjects from Alzheimer’s Disease Neuroimaging Initiative and demonstrate that it can significantly reduce variability in streamline density, improve consistency in along-tract microstructure profiles, and provide useful metrics for automated bundle quality control. These results suggest that SDNorm can help enhance the reproducibility and robustness of bundle reconstruction across heterogeneous image acquisition protocols and tractography settings, making it well-suited for large-scale and multi-site neuroimaging studies.

## 1 Introduction

Tractometry is an approach that map scalar microstructural measures along the brain’s white matter (WM) fiber bundles, which can be identified by tractography. Tractometry analysis heavily depends on the consistency and accuracy of tractography and bundle segmentation. However, both steps suffer from high variability across subjects, anatomical bundles, and diffusion MRI (dMRI) acquisitions.

Tractography requires the selection of numerous parameters—such as step size, seeding strategy, angle threshold and stopping criteria—whose optimal values may differ across studies. Bundle segmentation methods also rely on modifiable parameters such as distance thresholds, inclusion/exclusion criteria for ROI definitions, and atlas alignment. Even when processing parameters are held constant, anatomical differences across individuals and variability in data quality can affect the ability to reconstruct bundles of interest [16]. The inconsistency in these processing parameters directly impacts how scalar maps are sampled and interpolated in tractometry analyses. For example, regions with low streamline density may produce noisier tract profiles. This variability in streamline representation can complicate group-level comparisons, which typically do not account for differences in spatial sampling. Therefore, reducing arbitrary sources of variability in tractography and bundle segmentation is essential, to improve the reliability and reproducibility of tractometry studies.

A common class of streamline filtering methods focuses on improving the biological validity of tractography by aligning streamlines with the underlying diffusion signals or estimated fiber orientation distributions. Notable examples include Spherical-Deconvolution Informed Filtering of Tractograms (SIFT and SIFT2) [18], [19], Convex Optimization Modeling of Microstructure Informed Tractograpy (COMMIT) [4], and Linear Fascicle Evaluation (LiFE) [13]. These supervised filtering methods are primarily designed for whole-brain tractograms in the context of connectivity analysis. However, they are not applicable to individual white matter (WM) bundles, as multiple tracts often intersect within the same voxel, or even fixel [17] at the current spatial resolution of diffusion MRI. In contrast, bundle-specific filtering methods —such as FiberNeat[3] and BundleCleaner [5]—remove anatomically implausible streamlines from bundles in an unsupervised approach, as a form of quality control. They assume coherent streamline trajectories and use clustering algorithms to remove streamlines. While these methods are effective when a bundle is overestimated—i.e., adequately reconstructed but contains false positive streamlines, they have limited use when a bundle is underestimated—i.e., has sparse streamlines and fails to capture the bundle’s full spatial extent. This asymmetry in handling false positives and false negatives presents a key challenge for bundle-level quality control. Applying filtering indiscriminately to all extracted bundles may discard valid but sparse reconstructed fibers. Ideally, a robust approach should identify underestimated bundles while selectively filtering only overestimated bundles.

In this study, we introduce **Streamline Density Normalization (SD-Norm)**, a two-step procedure that aims to reduce arbitrary sources of variability in extracted bundles across subjects and tractography settings. We propose to *normalize* the streamline density of bundles using a supervised approach instead of *filtering* to remove false positive streamlines. SDNorm first computes streamline weights to match an individual bundle to a template density map derived from a reference group using ridge regression. We also introduce a novel metric called **effective streamline point density (eSPD)** that describes the average number of streamline points traversing a unit voxel for a given bundle. In the second step of SDNorm, we iteratively prune each bundle based on streamline weights, until a target eSPD is reached. We assess SDNorm on multi-site diffusion MRI from the Alzheimer’s Disease Neuroimaging Initiative (ADNI), and show that SDNorm can significantly reduce variability in streamline density and tractometry profiles across subjects and tractography settings. Moreover, metrics describing the model fit of the ridge regression model offer utility for automated QC and can help flag bundles that are underestimated or poorly constructed. Code for SDNorm is available at https://github.com/wendyfyx/SDNorm.

## 2 Data & Materials

### 2.1 Diffusion MRI Processing

To create the template density maps, we processed dMRI data from 200 sex-matched subjects (100 male / 100 female; 36-100 years old; mean age: 58.75 *±* 14.86 (SD) years) from the Lifespan Human Connectome Project Aging (HCP-A) study[9] using processing steps described in [6]. All preprocessed DWI volumes have 1.5-mm isotropic voxel size. Fiber orientations were reconstructed using multi-shell multi-tissue constrained spherical deconvolution (msmtCSD) [10].

We analyzed DWI data from 140 participants from ADNI3 (**Table 1**) [24]. For each imaging protocol, we selected 20 subjects matched by sex. Within each protocol group, the cohort was further balanced by cognitive status, consisting of 10 cognitively normal (CN) individuals and 10 individuals diagnosed with either mild cognitive impairment (MCI) or dementia. Additional information on acquisition protocols and DWI preprocessing steps is available in [7][21]. All DWI were resampled to 2-mm isotropic voxel size; fiber orientations were reconstructed using CSD[22] for single-shell diffusion protocols, except for S127, where msmtCSD was used.

**Table 1.** ADNI3 participant characteristics by protocol (N=140). Protocol abbreviations are prefixed with the scanner vendor, where S stands for Siemens, P for Philips, and GE for General Electric. The number refers to the number of diffusion-weighted gradients. All protocols are single-shell with b=1000 *s/mm*^2^, except for S127, that is multi-shell.

| Protocol | Sex | Age | Diagnosis |  |  |
| --- | --- | --- | --- | --- | --- |
|  |  |  | CN | MCI | Dementia |
| GE36 | 10 M / 10 F | $71.5 \pm 6.8$ | 10 | 9 | 1 |
| GE54 | 10 M / 10 F | $78.7 \pm 7.7$ | 10 | 7 | 3 |
| P33 | 10 M / 10 F | $76.8 \pm 8.7$ | 10 | 6 | 4 |
| P36 | 10 M / 10 F | $72.0 \pm 7.0$ | 10 | 9 | 1 |
| S127 | 10 M / 10 F | $72.7 \pm 7.4$ | 10 | 8 | 2 |
| S31 | 10 M / 10 F | $72.8 \pm 7.5$ | 10 | 9 | 1 |
| S55 | 10 M / 10 F | $75.7 \pm 9.0$ | 10 | 6 | 4 |

### 2.2 Tractography Processing

For all HCP-A and ADNI3 subjects, we performed bundle-based tractography to extract 10 bundles - the arcuate fasciculus left (AF_L) and right (AF_R), cingulum frontal parietal left (C_FP_L) and right (C_FP_R), corpus callosum forceps major (CC_ForcepsMajor) and forceps minor (CC_ForcepsMinor), corticospinal tract left (CST_L) and right (CST_R), inferior fronto-occipital fasciculus left (IFOF_L) and right (IFOF_R). Inspired by Bundle-Specific Tractography [15] and Automatic Fiber Tracking from DSI-Studio [25], seeds were only set within the bundles of interest to constrain fiber tracking.

Atlas bundles were obtained from the population-averaged HCP-1065 Young Adult atlas [26]. To obtain the transform between the subject space and MNI space, a fractional anisotropy (FA) map was first computed for each ADNI3 subject by fitting DTI, and registered to the ICBM 2009a Nonlinear Asymmetric T1w template (1-mm isotropic voxel size) using ANTS Syn [23]. To create the seed mask, each atlas bundle was converted into a binary mask, dilated, and warped to the subject’s dMRI space with the ANTS’ non-linear transform. Eight seeds were randomly placed for each voxel within the seed mask. A binary stopping criterion was defined as the intersection of the dilated seed mask and FA *>* 0.05. We selected a relatively low FA threshold to ensure the streamlines are adequately reconstructed in regions with potentially reduced anisotropy due to neurodegeneration in ADNI3 subjects. For each of the 10 bundles of interest, local probabilistic tracking from DIPY [8],[1] was used to generate streamlines with maximum angle of 20^°^ and step size of 0.5 mm. Streamlines were retained if their lengths fell within the minimum and maximum range defined by the atlas bundle, with a 20% tolerance margin. After fiber tracking, the atlas bundles were warped to subject space and used to filter the generated streamlines with the Fast Streamline Search (FSS) algorithm [20]. FSS was chosen for its computational efficiency and simplicity, requiring only a single parameter —the search radius. We used a radius of 6 mm for all bundles, except for the IFOF, where a radius of 7 mm provided better segmentation results.

To evaluate the performance of SDNorm across tractography settings in ADNI3 subjects, we ran bundle-based tractography with the following modification to the parameters: the number of seeds per voxel was increased to 12, and the step size was decreased to 0.2 mm. We refer to this configuration as *Run B*, while the initial configuration with 8 seeds and 0.5 mm step size is referred to as *Run A*.

All bundles from all subjects and runs were manually quality-checked (QC) and labeled as “Pass,” “Poor,” or “Fail” based on their anatomical plausibility and streamline coverage. Bundles labeled “Poor” typically have limited coverage or contain visible false positives but could still be usable with additional processing. “Fail” labels were assigned to bundles with severe reconstruction issues—such as very few streamlines, large gaps, or anatomically implausible trajectories. All bundles from HCP-A subjects passed QC for template creation. Details for ADNI3 bundle QC are in **Section 4.1**.

## 3 Methods

In SDNorm, template density maps are first created for each bundle of interest from a reference group. Streamline weights are then calculated to match each subject bundle’s density map to match the template, followed by a pruning procedure. We describe each step in detail below.

### 3.1 Template Map Creation

A density map is defined as the number of streamlines passing through voxels in a reference volume. Template density maps from 10 bundles of interest were computed from 200 HCP-A subjects (see **Figure 1(a))**. As the HCP pipeline produces processed dMRI in the MNI space, there was no need for registration to align density maps across subjects. For each bundle density map, voxels where only one streamline passes through were removed, and the map was normalized so that all values sum to 1. The normalized density maps were then averaged across subjects for each bundle. Gaussian smoothing (*σ* = 0.5) and log-filtering (voxels with the top 50% percentile log-density were retained) were then applied to the group average density map [11] to create the final template map.

**Fig. 1.**
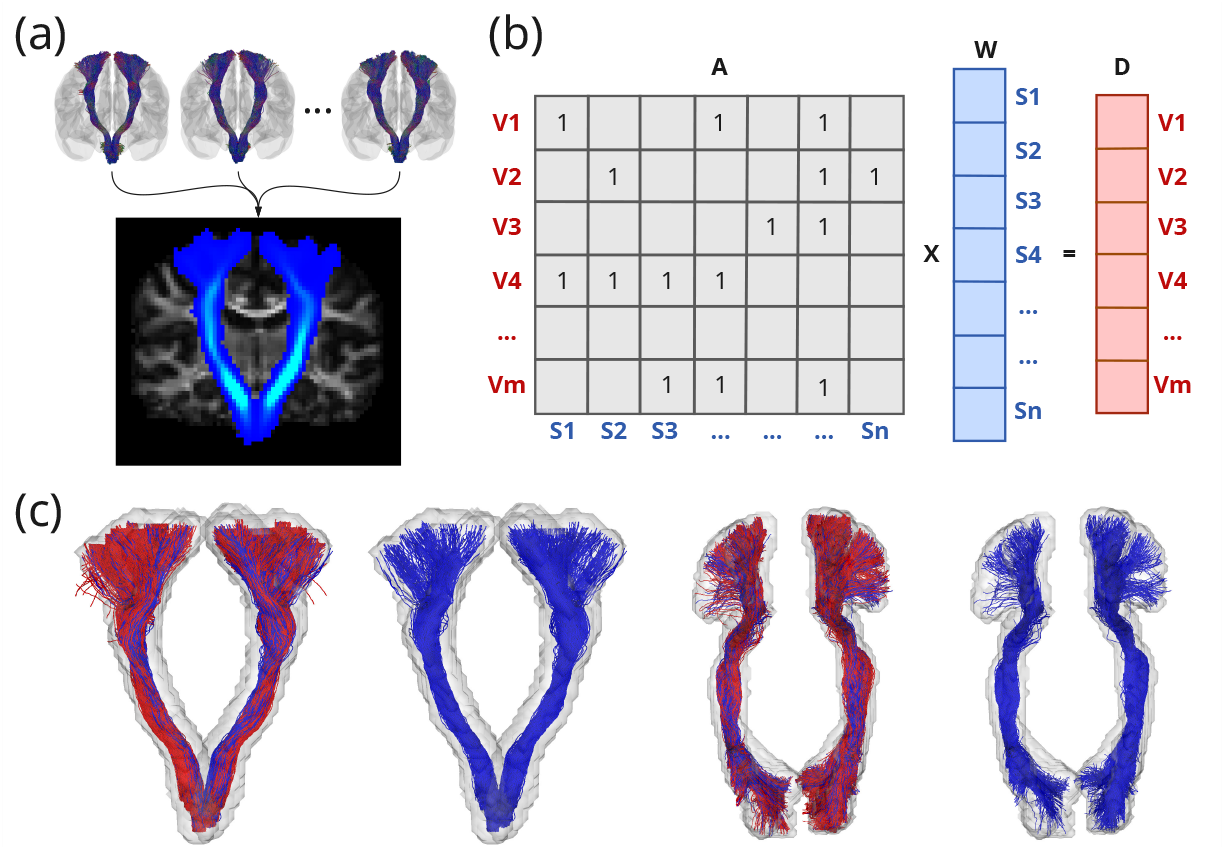
Streamline Density Normalization (SDNorm). **(a)** Creating the group-level streamline density map of the corticospinal tract (CST) from a reference group. **(b)** SDNorm Step 1: Calculating streamline weights. **c** Examples bundles before and after SDNorm Step 2 pruning. *red* : streamlines discarded; *blue*: remaining streamlines.

### 3.2 SDNorm Step 1: Streamline Weight Calculation

The goal of this step is to assign one weight per streamline such that the weighted streamline density map for a given bundle best matches the corresponding template map, subject to regularization (see **Figure 1(b))**. We create the streamline-voxel adjacency matrix *A* ∈ ℝ^*V ×S*^, where each column *A*_*i*_ is a binary vector indicating which voxels streamline *i* passes through, *V* is the number of voxels with non-zero streamline density in the reference volume, and *S* is the total number of streamlines in a given bundle. We compute weights *w* ∈ ℝ^*S*^ with the following objective:

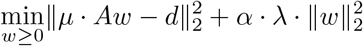

where *d* ∈ ℝ^*V*^ is the flattened template density map and *λ* is the ridge regularization weight. The first term matches streamline density map to the template, and the second term regularizes the weights to prevent overfitting. Inspired by the scaling parameters in SIFT2 [19], we defined *µ*, a fixed parameter to match the scale between streamline density and the normalized template map, and *α*, another fixed scaling parameter to control the effect of regularization as

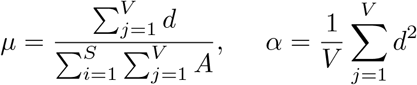

Both *µ* and *α* are calculated prior to optimization, and *w* is the only free parameter.

### 3.3 SDNorm Step 2: Streamline Pruning

After the streamline weights *w* are computed, they can be used to weight streamline contributions in the second pruning procedure. Although the true underlying bundle-specific fiber density may differ, we aim to reduce inter-bundle density variability using the following pruning procedure: For a given bundle, we define the metric Streamline Point Density (SPD) as the total number of streamline points for a given bundle divided by the bundle volume. Suppose 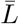 is the average streamline length and *c* is the step size, both defined in millimeters (mm), the average number of points per streamline is 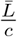. We can instead compute

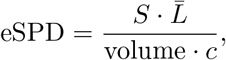

or **Effective Streamline Point Density (eSPD)**. This formulation makes it easier to prune streamlines: given a target eSPD, the number of streamlines to retain at iteration *k* is

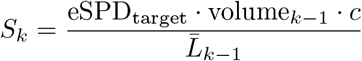

At each iteration, *S*_*k*_ streamlines with the highest weights in *w* are retained, and volume and 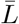 are recomputed. We iteratively prune the bundle using this procedure until the desired eSPD is reached.

### 3.8 Experiments

In this study, we applied SDNorm to both run A and B bundles from 140 ADNI3 subjects. For each bundle, the corresponding template map is first warped to the subject space using the ANTS transform computed in **Section 2.2**. Due to difference in resolution between the template maps (1-mm isotropic) and subject space FA (2-mm isotropic), SDNorm step 1 is much faster when *A* and *d* are defined in the subject space. All subject bundles from run B are resampled to *s* = 0.5 mm fixed step size to match run A. SDNorm pruning was applied with a target eSPD of 8. For bundles with more than 10,000 streamlines, QuickBundles was used to downsample bundles to following the approach in BundleCleaner [5], and speed up model fitting in SDNorm step 1.

To evaluate the effect of SDNorm on along-tract profiles, we computed mean FA profiles for each bundle using Bundle Analytics (BUAN) [2]. BUAN creates 100 along-tract segments, and the FA values projected on streamline points within each segment are averaged to create the mean FA profile.

## 4 Results

### 4.1 Model Fit & Quality Control

After calculating streamline weights in SDNorm step 1, we evaluate model fit using *R*^2^, the coefficient of determination. The average *R*^2^ values for bundles from run A and run B were 0.79 *±* 0.16 and 0.78 *±* 0.17 respectively. We show bundles with the highest and lowest *R*^2^ scores for 4 bundles in **Figure 2(b)**. Bundles with high *R*^2^ are more densely packed and have good streamline coverage, whereas bundles with low *R*^2^ are typically underestimated or contain streamlines with anatomically implausible trajectories. As *R*^2^ is a metric describing the fit between a weighted streamline density map and the template map, we also calculate the normalized cross correlation (NCC) between the original unweighted streamline density map and the template map. We plot *R*^2^ against NCC in **Figure 2(a)**, colored by labels from manual QC. Bundles labeled either “Poor” or “Fail” are concentrated in the lower left quadrant. Using thresholds of *R*^2^ = 0.4 or *NCC* = 0.5 can well distinguish bundles that are labeled “Pass” (with precision of 0.985 and 0.983, and recall of 0.991 and 0.989 respectively) and both metrics show promise in automated bundle QC.

**Fig. 2.**
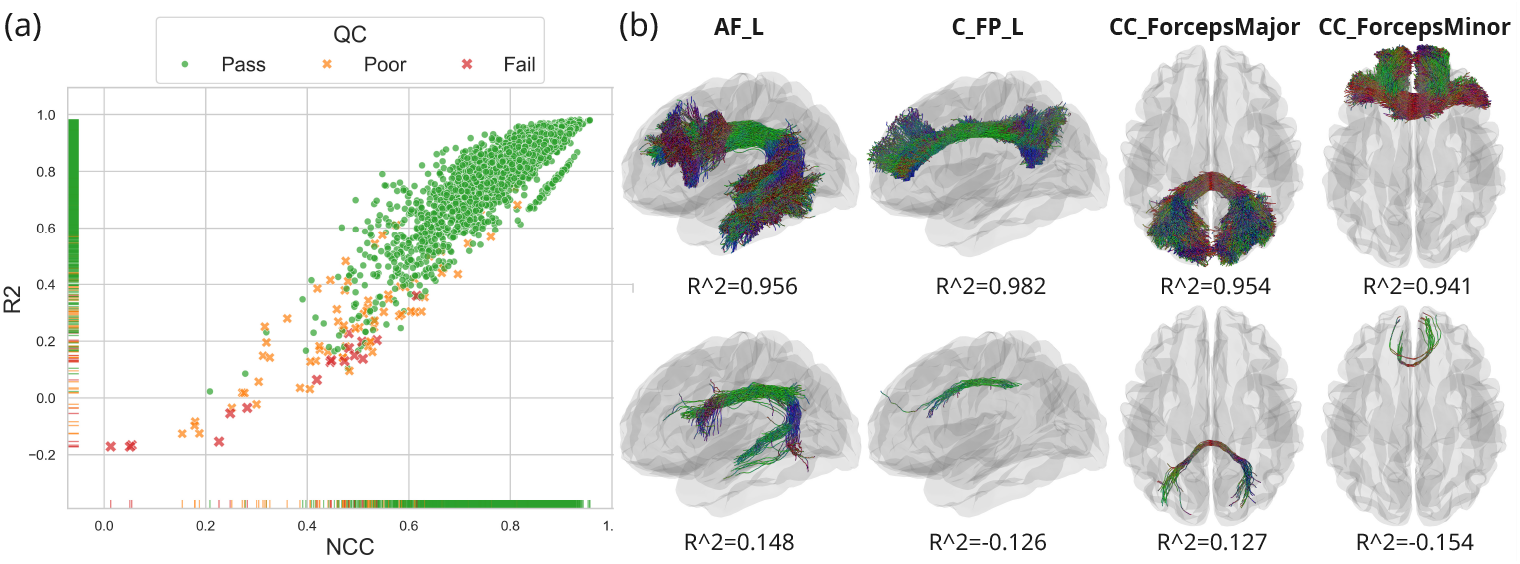
Quality Control (QC) using SDNorm. **(a)** Normalized Cross Correlation (NCC) between the subjects’ density maps and template map is plotted against *R*^2^ from SDNorm Step 1, colored by QC labels. **(b)** Subject bundles with the highest and lowest *R*^2^ across 4 bundles.

### 4.2 Evaluation on Streamline Density

To evaluate the effectiveness of SDNorm in reducing variability across tractography settings, we compared the streamline density distributions between Run A and Run B using Quantile-Quantile (Q-Q) plots for all ADNI3 subjects, stratified by acquisition protocol. Results for three representative bundles are shown in **Figure 3**. The black dashed line represents the reference distribution where both runs have identical density distributions. Prior to SDNorm, the Q-Q plots for the original bundles (red) exhibit consistently steep slopes, indicating that Run B bundles have substantially higher streamline densities than Run A—a result expected due to the increased seeding density used in Run B. After applying SDNorm, the Q-Q plots (blue) align much more closely with the reference line, indicating that the density distribution are much more similar between two runs. Aside from the change in slopes in the Q-Q plots, SDNorm also suppresses extreme values in the density distribution. For instance, peak densities reached nearly 2,000 streamlines per voxel in some CC_ForcepsMajor bundles before SDNorm, whereas peak density values were consistently reduced to below 300 streamlines per voxel after SDNorm.

**Fig. 3.**
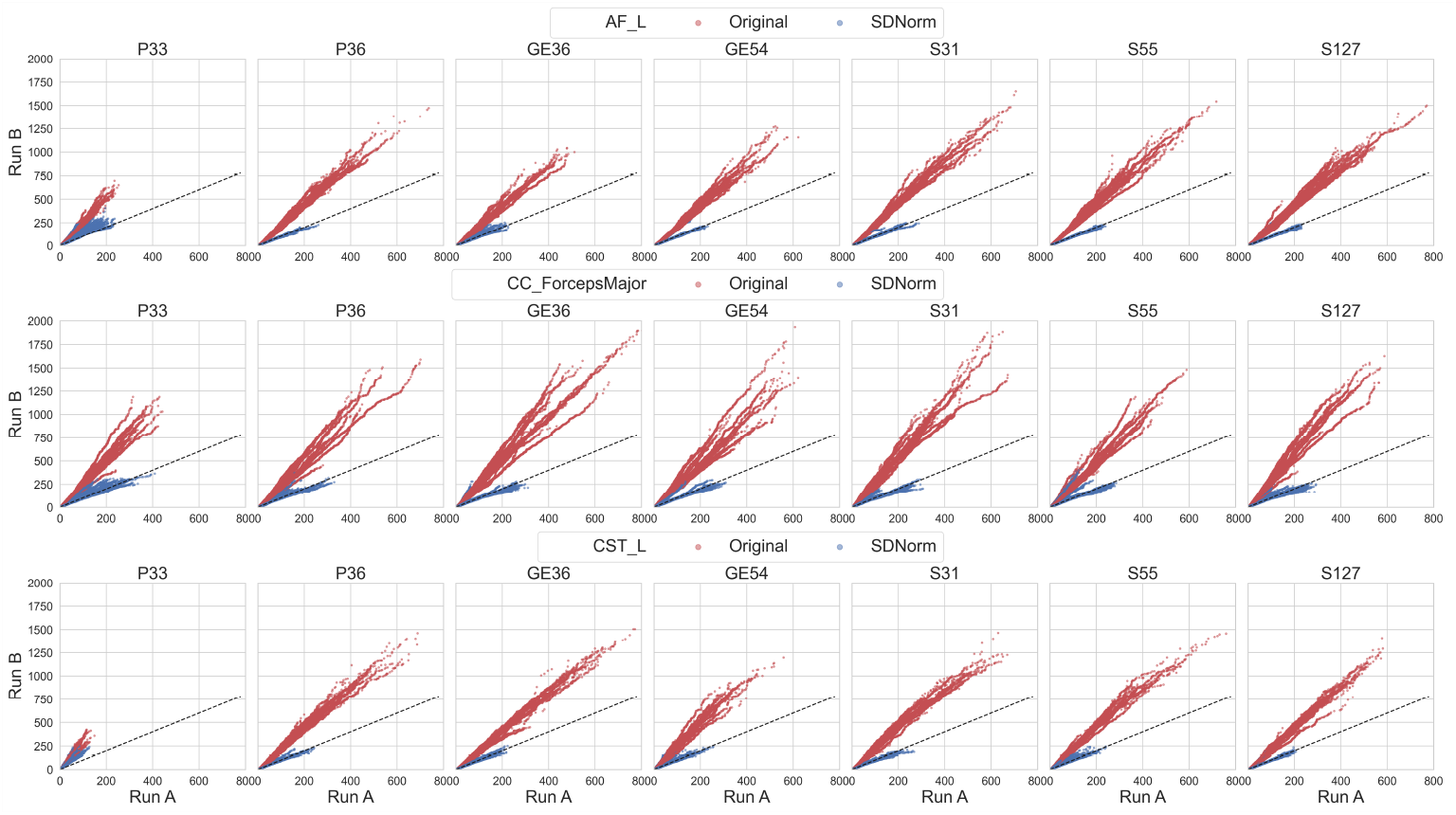
Quantile-Quantile (Q-Q) Plot of density maps values from Run A bundles again those from Run B bundles before and after applying SDNorm, stratified by acquisition protocol across 3 bundles.

We also observed moderate variability in streamline density across different bundles and acquisition protocols. Prior to SDNorm, CC_ForcepsMajor exhibited both higher peak densities and greater inter-subject variability compared to AF_L and CST_L. Protocol P33, in particular, yielded bundles with lower peak density than other protocols across bundles. CST_L bundles from P33 peak below 400 streamlines per voxel, and they likely had low eSPD values and were therefore not subject to pruning. Overall, these results demonstrate that SDNorm effectively reduces variability in streamline density across tractography settings, subjects and protocols.

### 4.3 Along-Tract Profile of Fractional Anisotropy

We further evaluated the impact of SDNorm on along-tract FA profiles by evaluating the similarity of profiles between Run A and Run B. For each bundle, we computed the L2 norm between the mean FA profiles from two runs, defined as FA_diff_ = ∥FA_*A*_ − FA_*B*_∥_2_, where FA_*A*_ and FA_*B*_ are the mean FA profiles defined along 100 segments for run A and B, as detailed in **Section 3.4**.

Since bundles from both runs are derived from the same diffusion data and the same FA map is sampled, we expect their mean FA profiles to be similar. However, differences in streamline geometry and density between the two tractography settings can introduce variability in how the scalar values are sampled and interpolated. To assess whether SDNorm reduces this variability, we performed a paired statistical analysis using the Wilcoxon signed-rank test, testing the hypothesis that FA_diff_ is significantly lower after SDNorm. **Figure 4** shows boxplots of FA_diff_ for all 10 bundles before and after applying SDNorm, along with the *p*-values from the Wilcoxon test. Across all bundles, the differences in mean FA profiles between Run A and Run B were significantly reduced following SDNorm (*p* < 0.05, FDR corrected), indicating improved consistency in FA profiles. Among the bundles evaluated, CC_ForcepsMajor exhibited the largest FA_diff_ values and the greatest inter-subject variance, both before and after SD-Norm. This is consistent with their higher streamline density and geometric variability observed in earlier analyses.

**Fig. 4.**
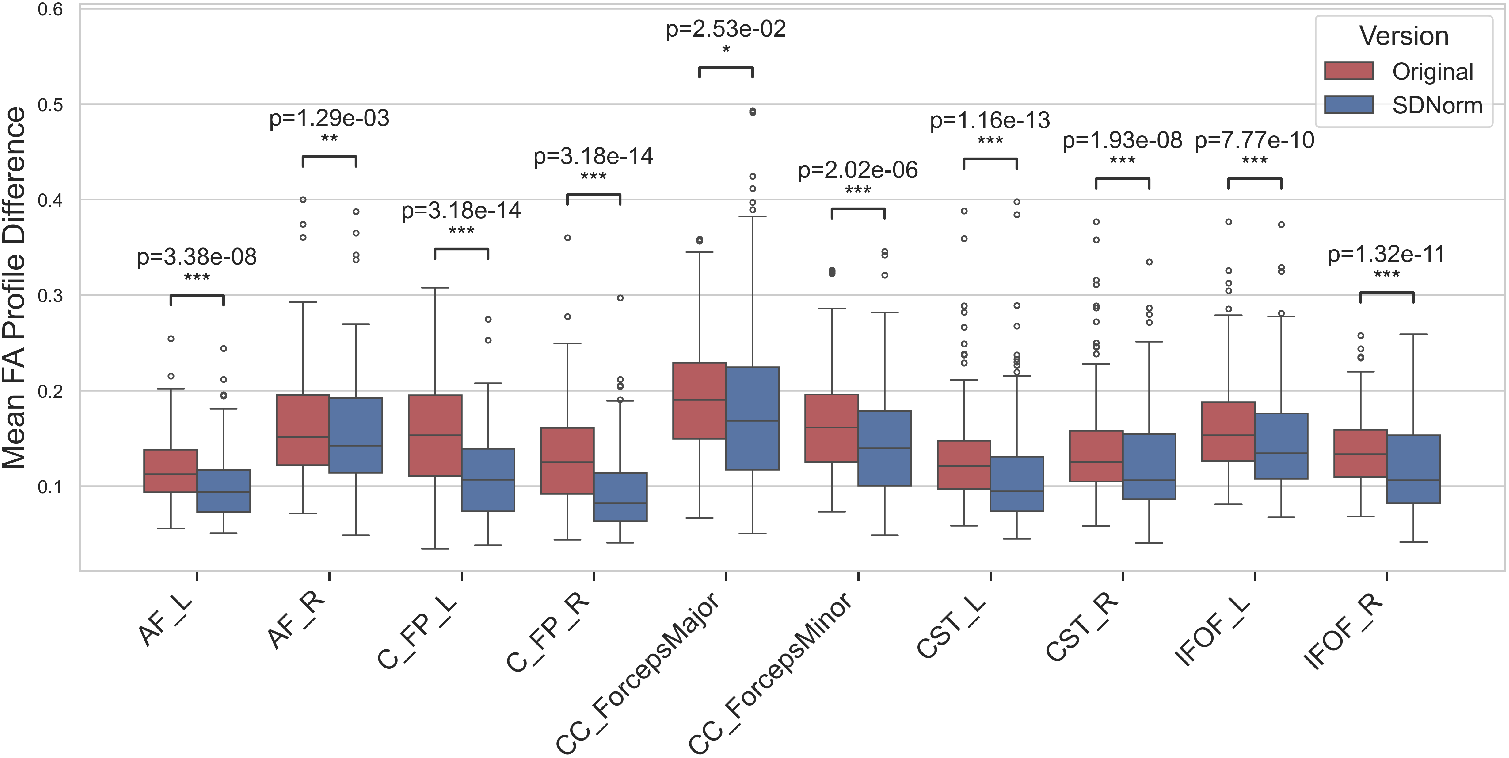
Boxplots of FA_diff_ before and after applying SDNorm across 10 bundles. Significance levels and FDR-corrected *p*-values from the Wilcoxon test is shown for each bundles (*: *p <* 0.05, **: *p <* 0.01, ***: *p <* 0.001).

These findings indicated that SDNorm leads to more stable and reproducible along-tract microstructural profiles, which is essential for downstream group analyses and clinical interpretation.

## 5 Discussion

In this study, we introduced SDNorm, a supervised method for normalizing bundle streamline density across tractography settings and subjects. SDNorm is a two step procedure, and contains only 3 user-defined parameters—step size, regularization weight, and target eSPD.

A potential concern is that SDNorm may reduce subject-level variability when matching subject bundles to a template map. Importantly, SDNorm does not “upsample” bundles with streamline densities below the target eSPD threshold, thereby preserving sparse reconstructions that may reflect biological variability. Users may select higher eSPD thresholds to retain more subject variability and lower thresholds to apply stricter density normalization. In tractometry applications, statistical stability of microstructural profiles can vary across datasets and bundles of interest, as their values depend on acquisition protocols[14]. In this study, we have shown that FA profiles are more reproducible across tractography settings after applying SDNorm. In future work, we aim to systematically investigate how different pruning thresholds may influence the stability of along-tract microstructural measures and the detection of biologically meaningful variability across subjects.

An important consideration when using SDNorm is the choice of template density maps. We recommend that users generate a study-specific template map using a subset of their study cohort or a demographically matched reference group processed with similar bundle extraction pipelines. To support this, we provide code for constructing the template maps in the SDNorm Github repository.

Lastly, we note that SDNorm was developed and evaluated using bundles generated with probabilistic tractography. Deterministic tractography is less sensitive to uncertainty in fiber orientations and tends to generate streamlines that are more densely packed in the core of WM bundles, while probabilistic tractography tends to produce a wider spread of streamlines [12]. Due to such differences, we do not recommend using templates generated from deterministic tractography when normalizing bundles from probabilistic tractography, or *vice versa*. Future work will investigate how SDNorm performs on deterministic tractography.

## 6 Conclusion

We propose SDNorm, a supervised method for reducing bundle variability by normalizing streamline density. By incorporating a model fitting step and the novel resolution-independent eSPD metric in pruning, SDNorm requires three intuitive user-defined parameters and can be applied to different anatomical bundles. Applied to multi-site data from ADNI3, we show that SDNorm can reduce differences in streamline density distribution across subjects and tractography settings and improve the reproducibility of along-tract FA profiles.

In addition, model fit metrics derived from SDNorm can assist with automated quality control by identifying poorly reconstructed bundles. This approach offers a practical and scalable solution for improving the reliability of bundle extraction and tractometry in studies with multi-site dMRI data.

## Acknowledgments

This study was supported by the National Institute on Aging (NIA) under grant RF1-NS136995, the National Institute of Health (NIH) under grant NIH-T32AG058507, National Institute of Mental Health under grant R01MH134004 and the US Alzheimer’s Association under grant AARG-23-1149996.

## References

1. Behrens, T., et al.: Characterization and propagation of uncertainty in diffusion-weighted MR imaging. Magnetic Resonance in Medicine 50(5), 1077–1088 (Nov 2003). 10.1002/mrm.10609

2. Chandio, B.Q., et al.: Bundle analytics, a computational framework for investigating the shapes and profiles of brain pathways across populations. Scientific Reports 10(1), 17149 (Dec 2020). 10.1038/s41598-020-74054-4

3. Chandio, B.Q., et al.: FiberNeat: unsupervised streamline clustering and white matter tract filtering in latent space. preprint, Neuroscience (Oct 2021). 10.1101/2021.10.26.465991

4. Daducci, A., et al.: COMMIT: Convex Optimization Modeling for Microstructure Informed Tractography. IEEE Transactions on Medical Imaging 34(1), 246–257 (Jan 2015). 10.1109/TMI.2014.2352414

5. Feng, Y., et al.: BundleCleaner: Unsupervised Denoising and Subsampling of Diffusion MRI-Derived Tractography Data. In: Computational Diffusion MRI, vol. 14328, pp. 152–164. Springer Nature Switzerland, Cham (2023). 10.1007/978-3-031-47292-3_14

6. Feng, Y., et al.: BundleAGE: Predicting White Matter Age using Along-Tract Microstructural Profiles from Diffusion MRI. In: 2024 20th International Symposium on Medical Information Processing and Analysis (SIPAIM). IEEE, Antigua, Guatemala (Nov 2024). 10.1109/SIPAIM62974.2024.10783507

7. Feng, Y., et al.: Microstructural mapping of neural pathways in Alzheimer’s disease using macrostructure-informed normative tractometry. Alzheimer’s & Dementia p. alz.14371 (Dec 2024). 10.1002/alz.14371

8. Garyfallidis, E., et al.: Dipy, a library for the analysis of diffusion MRI data. Frontiers in Neuroinformatics 8 (Feb 2014). 10.3389/fninf.2014.00008

9. Harms, M.P., et al.: Extending the Human Connectome Project across ages: Imaging protocols for the Lifespan Development and Aging projects. NeuroImage 183, 972–984 (Dec 2018). 10.1016/j.neuroimage.2018.09.060

10. Jeurissen, B., et al.: Multi-tissue constrained spherical deconvolution for improved analysis of multi-shell diffusion MRI data. NeuroImage 103, 411–426 (Dec 2014). 10.1016/j.neuroimage.2014.07.061

11. Maffei, C., et al.: Insights from the IronTract challenge: Optimal methods for mapping brain pathways from multi-shell diffusion MRI. NeuroImage 257, 119327 (Aug 2022). 10.1016/j.neuroimage.2022.119327

12. Maier-Hein, K.H., et al.: The challenge of mapping the human connectome based on diffusion tractography. Nature Communications 8(1), 1349 (Nov 2017). 10.1038/s41467-017-01285-x

13. Pestilli, F., et al.: Evaluation and statistical inference for human connectomes. Nature Methods 11(10), 1058–1063 (Oct 2014). 10.1038/nmeth.3098, number: 10 Publisher: Nature Publishing Group

14. Reid, L.B., Cespedes, M.I., Pannek, K.: How many streamlines are required for reliable probabilistic tractography? Solutions for microstructural measurements and neurosurgical planning. NeuroImage 211, 116646 (May 2020). 10.1016/j.neuroimage.2020.116646

15. Rheault, F., et al.: Bundle-specific tractography with incorporated anatomical and orientational priors. NeuroImage 186, 382–398 (Feb 2019). 10.1016/j.neuroimage.2018.11.018

16. Schilling, K.G., et al.: Fiber tractography bundle segmentation depends on scanner effects, vendor effects, acquisition resolution, diffusion sampling scheme, diffusion sensitization, and bundle segmentation workflow. NeuroImage 242, 118451 (Nov 2021). 10.1016/j.neuroimage.2021.118451

17. Schilling, K.G., et al.: Prevalence of white matter pathways coming into a single white matter voxel orientation: The bottleneck issue in tractography. Human Brain Mapping 43(4), 1196–1213 (Mar 2022). 10.1002/hbm.25697

18. Smith, R.E., et al.: SIFT: Spherical-deconvolution informed filtering of tractograms. NeuroImage 67, 298–312 (Feb 2013). 10.1016/j.neuroimage.2012.11.049

19. Smith, R.E., et al.: SIFT2: Enabling dense quantitative assessment of brain white matter connectivity using streamlines tractography. NeuroImage 119, 338–351 (Oct 2015). 10.1016/j.neuroimage.2015.06.092

20. St-Onge, E., et al.: Fast Streamline Search: An Exact Technique for Diffusion MRI Tractography. Neuroinformatics 20(4), 1093–1104 (Oct 2022). 10.1007/s12021-022-09590-7

21. Thomopoulos, S.I., et al.: Diffusion MRI metrics and their relation to dementia severity: effects of harmonization approaches. In: 17th International Symposium on Medical Information Processing and Analysis. p. 79. SPIE, Campinas, Brazil (Dec 2021). 10.1117/12.2606337

22. Tournier, J.D., et al.: Robust determination of the fibre orientation distribution in diffusion MRI: non-negativity constrained super-resolved spherical deconvolution. NeuroImage 35(4), 1459–1472 (May 2007). 10.1016/j.neuroimage.2007.02.016

23. Tustison, N.J., et al.: The ANTsX ecosystem for quantitative biological and medical imaging. Scientific Reports 11(1), 9068 (Apr 2021). 10.1038/s41598-021-87564-6

24. Weiner, M.W., et al.: The Alzheimer’s Disease Neuroimaging Initiative 3: Continued innovation for clinical trial improvement. Alzheimer’s & Dementia: The Journal of the Alzheimer’s Association 13(5), 561–571 (May 2017). 10.1016/j.jalz.2016.10.006

25. Yeh, F.C.: Shape analysis of the human association pathways. NeuroImage 223, 117329 (Dec 2020). 10.1016/j.neuroimage.2020.117329

26. Yeh, F.C.: Population-based tract-to-region connectome of the human brain and its hierarchical topology. Nature Communications 13(1), 4933 (Aug 2022). 10.1038/s41467-022-32595-4

